# Empowering medaka fish biology with versatile genomic resources in MedakaBase

**DOI:** 10.1101/2025.05.13.653297

**Authors:** Kenji Morikami, Yasuhiro Tanizawa, Masaru Yagura, Mika Sakamoto, Shoko Kawamoto, Yasukazu Nakamura, Katsushi Yamaguchi, Shuji Shigenobu, Kiyoshi Naruse, Satoshi Ansai, Shigehiro Kuraku

## Abstract

Medaka, a group of small, mostly freshwater fishes in the teleost order Beloniformes, includes the rice fish *Oryzias latipes*, which is a prominent model organism for diverse biological fields. Chromosome-scale genome sequences of the Hd-rR strain of this species were obtained in 2007, and its improved version has facilitated various genome-wide studies. However, despite its widespread utility, omics data for *O. latipes* are dispersed across various public databases and lack a centralized platform. To address this, the medaka section of the National Bioresource Project (NBRP) of Japan established a genome informatics team in 2022 tasked with providing versatile *in silico* solutions for bench biologists. This initiative led to the launch of MedakaBase (https://medakabase.nbrp.jp), a web server that enables gene-oriented analysis including exhaustive sequence similarity searches. MedakaBase also provides genome-wide browsing of diverse datasets, including tissue-specific transcriptomes and intraspecific genomic variations, integrated with gene models from different sources. Additionally, the platform offers gene models optimized for single-cell transcriptome analysis, which often requires coverage of the 3′ untranslated region (UTR) of transcripts. Currently, MedakaBase provides genome-wide data for seven *Oryzias* species, including original data for *O. mekongensis* and *O. luzonensis* produced by the NBRP team. This article outlines technical details behind the data provided by MedakaBase.

## 1. Introduction

The family Adrianichtyidae, commonly referred to as medaka or ricefishes, is a group of small, mostly freshwater teleost fishes that belong to the order Beloniformes. Medaka fishes are widely distributed throughout East and Southeast Asia^1,2^ and consist of ∼40 species that are classified in two genera *Oryzias* and *Adrianichtys*, with the latter phylogenetically nested within the former (Figure 1). The medaka (*Oryzias latipes*) is a well-established model vertebrate that has gained increasing prominence in developmental and evolutionary biology.^3-5^ While zebrafish (*Danio rerio*) has long been the dominant teleost model, medaka offers distinct advantages that complement and, in some contexts, surpass zebrafish in life science research.^4,6^ These advantages include a compact genome (approximately half the size of the zebrafish genome; see below for details), highly inbred strains with well-documented genetic diversity, and the ability to tolerate a broad temperature and osmolarity range during embryonic development. Unlike zebrafish, which requires relatively warm water conditions, medaka can thrive across a wider range of environmental conditions, making them particularly useful for studying ecological adaptations. Additionally, their resilience to inbreeding has facilitated the development of isogenic lines, enabling precise genetic analyses.^7^

**Figure 1.**
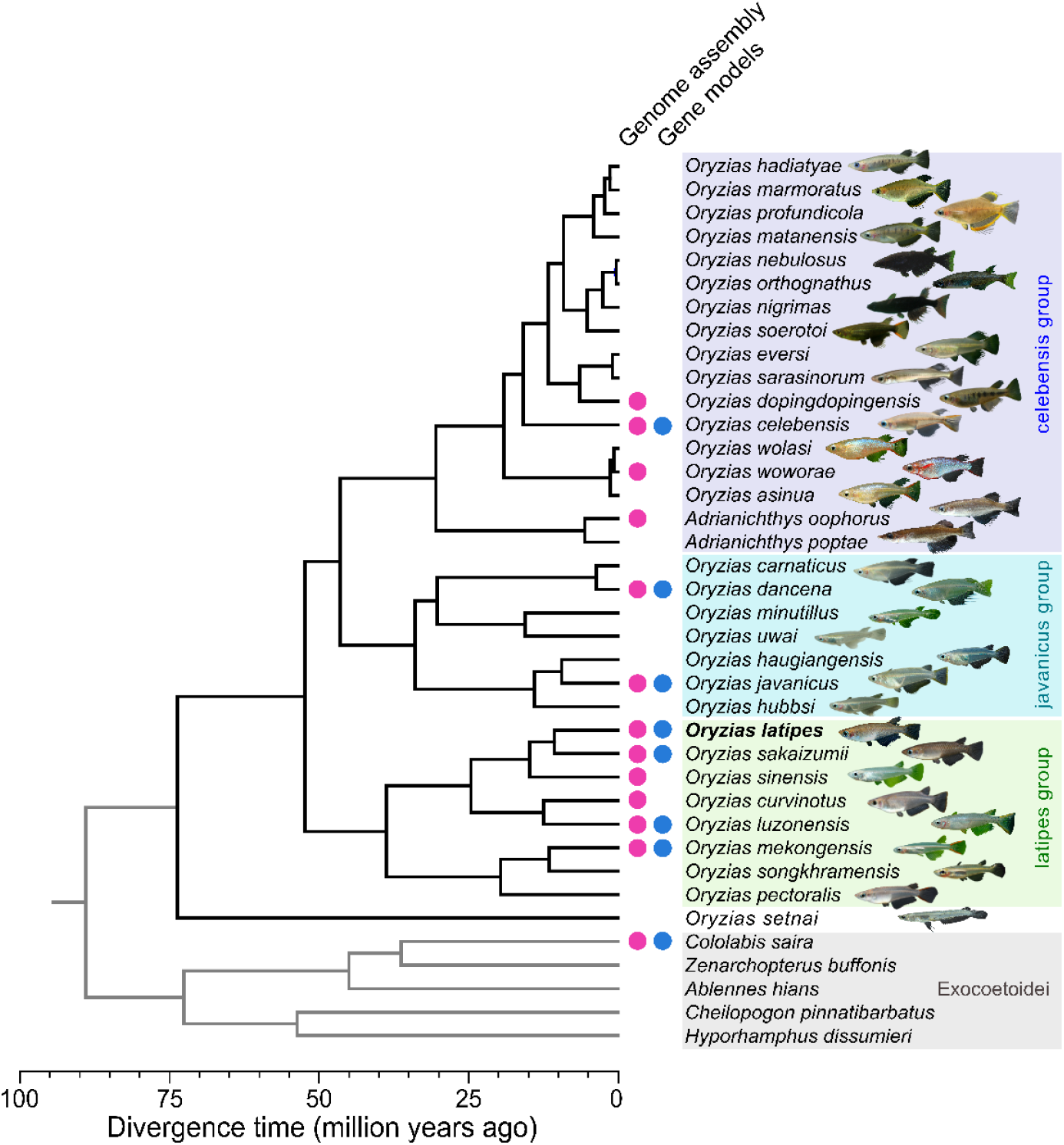
Phylogeny of the medaka fishes. Relationships and divergence times were inferred from molecular phylogenetic data.^52^ Five non-medaka species are included as outgroup. Magenta and blue circles indicate the species with available genome assemblies and gene models, respectively, available at NCBI^53^ (https://www.ncbi.nlm.nih.gov/datasets/genome/?taxon=76071) or other public repositories. The availability of the whole genome sequence data for *O. dancena* is based on that of *O. melastigma* which is now regarded as *O. dancena*.

Japan has played a central role in medaka research for over a century, particularly in genetics and sex determination studies. Medaka holds historical significance as the first species in which Y-linked inheritance was demonstrated^8^ and as the first vertebrate in which sex reversal was successfully induced.^9^ These foundational studies culminated in the identification of *Dmy*, the male-determining gene-the first non-mammalian equivalent of SRY.^10^ The medaka model has also contributed extensively to ecotoxicology and carcinogenesis research, serving as an important system for studying environmental and genetic factors affecting vertebrate development and disease.^11^ Unlike zebrafish, which lack heteromorphic sex chromosomes, medaka possesses a well-characterized XX/XY sex determination system, making it an ideal model for studying vertebrate sex chromosome evolution. The adoption of CRISPR/Cas9 genome-editing technology^12^ has further expanded the utility of medaka in functional genomics, enabling precise gene manipulation for studying developmental processes, behavior, and disease models. Additionally, large-scale mutagenesis projects have generated extensive mutant libraries with unique phenotypes, highlighting the value of multiple teleost genetic models in comparative biology.^13-15^

With its small but fully sequenced genome, extensive genetic and genomic resources, well-annotated sex determination system, and adaptability to various experimental conditions, medaka continues to be a vital teleost model for uncovering the fundamental principles in genetics, developmental biology, and evolutionary research. However, despite its widespread utility, omics data for medaka fishes are dispersed across various public databases, lacking a centralized platform for easy access and interpretation. In 2022, NBRP Medaka incorporated a new mission to disseminate genomic information into its activities involving organisms, as reported in this article.

## 2. Materials and methods

### 2.1. Web server implementation

The MedakaBase server was built as the MarpolBase genome database^16,17^ and Cats-I^18^ and consisted of five docker containers including WebApollo^19^ and SequenceServer.^20^ Its genome browser is managed by the JBrowse^21^ module in WebApollo, and sequence similarity search is performed under SequencerServer 2.0.0 in which NCBI BLAST 2.12.0+^22^ is implemented.

### 2.2. Genome sequencing and assembly

High molecular weight DNA was extracted from male tissues, from which digestive tracts were removed, using Qiagen Genomic-tip. An SMRT sequence library was constructed with an SMRTbell Express Template Prep Kit 2.0 (Pacific Biosciences, CA, USA) and sequenced in a single 8M SMRT cell on a PacBio Sequel IIe system (Pacific Biosciences). The sequencing outputs were processed to generate CCS to obtain HiFi sequence reads of a total of 38.2 Gb for *O. mekongensis* and 25.6 Gb for *O. luzonensis*. From these reads, adapter sequences were removed using the program HiFiAdapterFilt.^23^ The obtained HiFi sequence reads were assembled using the program hifiasm v0.16.1^24^ with default parameters, and the resultant primary assemblies were subjected to protein-coding gene annotation and data release.

### 2.3. Gene prediction

Gene models were inferred with the program Helixer v0.3.2^25^ using the model vertebrate_v0.3_m_0080.h5 on the *O. mekongensis* and *O. luzonensis* genome contigs obtained as described above.

### 2.4. UTR extension of existing *O. latipes* Hd-rR gene models

Paired-end reads from short-read RNA-seq data derived from various tissues (SRP044784) were aligned to the reference genome assembly (ASM223467v1) with hisat2 v2.2.1.^26^ The obtained BAM file was input into the execution of the program peaks2utr v1.1.2.^27^ The GTF file with enriched 3′ UTR information by peaks2utr was used in the processing of the scRNA-seq data (GSM4959937^28^) produced with Chromium (10X Genomics) by 10X Genomics Cell Ranger v7.1.0. The modified GTF/GFF files we provide were obtained using peaks2utr with the default max-distance parameter.

## 3. Results

### 3.1. Existing medaka genome resources

The most long-standing genome resource for medaka fish was established in the inbred strain Hd-rR from southern Japanese species (*O. latipes*). It was initially released in 2007 as a product of Sanger sequencing^29^ (reviewed in refs 30,31). This was replaced with an improved version produced with long-read sequencing with consensus long reads (CLR) obtained by the single-molecule real-time (SMRT) technology of Pacific Biosciences (PacBio).^32^ This long-read-based genome assembly on a chromosome scale is available as the ‘reference’ (formerly ‘representative’) at NCBI with the identifier ASM223467v1 (GCF_002234675.1), which achieved one of the best completeness and continuity among well-studied teleost fishes.

The same Japanese group produced the chromosome-scale genome assemblies of the other major inbred strains that belong to closely-related species of the southern Japanese species (commonly referred to as the *Oryzias latipes* species complex); HNI of the northern Japanese species (*O. sakaizumii*) and HSOK of the east Korean species^32^ (*Oryzias* sp.). These resources are also available at Ensembl run by EBI.^33^ Both NCBI and Ensembl provide the predicted sets of genes that consist mostly of protein-coding genes, which however harbor significant differences between these two sources, e.g., in the components other than protein-coding genes and the coverage of non-coding exons (i.e., number of genes with UTR; see below) as well as the number of predicted protein-coding genes (Table 1).

**Table 1.** Statistics of existing and modified gene models featuring metrics related to scRNA-seq analysis.

| Gene models | NCBI<br>GFF | Ensembl<br>GFF | NBRP<br>improved by<br>peaks2utr (v1) | NBRP<br>improved by<br>peaks2utr (v3) |
| --- | --- | --- | --- | --- |
| Number of genes | 26,658 | 23,622 | 23,622 | 26,658 |
| Number of protein-coding genes* | 22,071 | 23,600 | 23,600 | 22,071 |
| Number of protein-coding genes with 5' UTR | 21,415 | 12,656 | 12,656 | 21,415 |
| Number of protein-coding genes with 3' UTR | 21,545 | 12,404 | 19,129 | 21,703 |
| Median length of 5' UTR | 128 | 201 | 201 | 128 |
| Median length of 3' UTR | 677 | 1,692 | 1,356 | 777 |
| Reads mapped confidently to transcriptome<sup>#</sup> (%) | 72.3 | 61.3 | 68.9 | 73.8 |
| Fraction of conserved orthologs recognized as 'Complete' by BUSCO (%) | 98.52 | 95.91 | 95.91 | 98.52 |
| Fraction of conserved orthologs recognized as 'Fragmented' by BUSCO (%) | 0.03 | 0.99 | 0.99 | 0.03 |
| Fraction of conserved orthologs recognized as 'Missing' by BUSCO (%) | 1.46 | 3.10 | 3.10 | 1.46 |

\*Number of protein-coding genes excluding antibody-related protein-coding genes (e.g., V\_gene\_segments). Median length of 3' UTR and 5' UTR, gene expression metrics by Cell Ranger and BUSCO values were evaluated on gene models including only protein-coding genes. <sup>#</sup>Gene expression metrics using Cell Ranger. Note that these statistics are based on the interpretation of GFF files, even though no explicit lines defined as UTRs exist in publicly available data files. Modified gene models v1 were based on gene models retrieved
from the Ensembl GTF file. Modified gene models v3 were based on gene models retrieved from the NCBI GFF file.

Since its initiation, Ensembl has been a long-standing platform for genome browsing and cross-species comparisons, including ortholog searches based on prebuilt molecular phylogenetic data (see https://asia.ensembl.org/info/genome/compara/homology_method.html). On the other hand, NIH hosting NCBI has more recently introduced its branch gateway Comparative Genomic Resources^34^ (CGR; https://www.ncbi.nlm.nih.gov/comparative-genomics-resource/), which provides various comparative solutions including a Comparative Genome Viewer (CGV; https://www.ncbi.nlm.nih.gov/cgv) that can instantly create a genome-wide synteny view, for example, between medaka and zebrafish (Figure 2). To fuel medaka biology with *in silico* resources and analysis solutions, we introduced useful websites and databases, including those mentioned above, on the Medaka Omics Reference page (https://github.com/Squalomix/medaka-annex).

**Figure 2.**
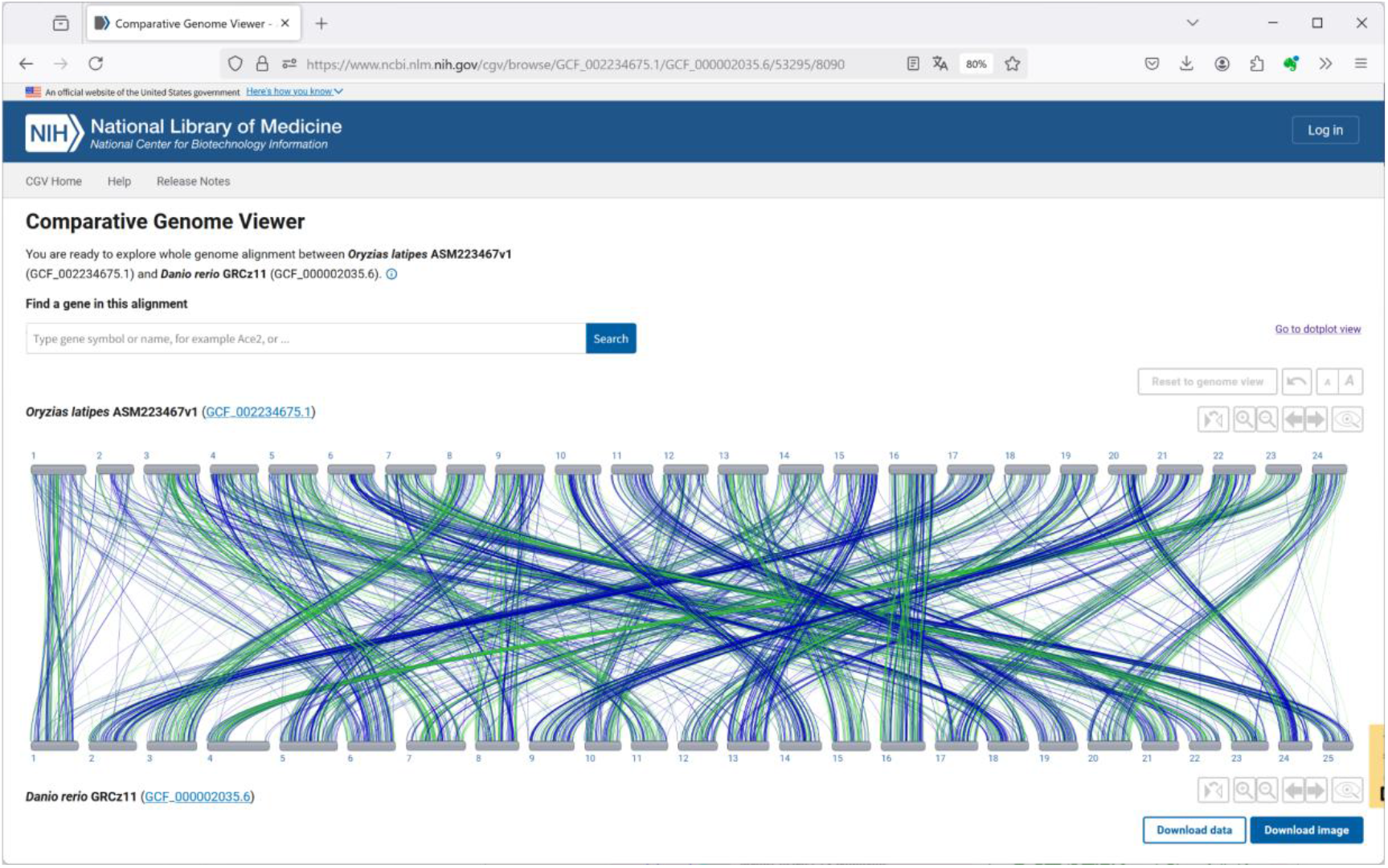
Genome-wide synteny conservation view between O. latipes Hd-rR and zebrafish. This graphic was produced in Comparative Genome Viewer (CGV) available at NIH, and is accessible under this URL (https://www.ncbi.nlm.nih.gov/cgv/browse/GCF_002234675.1/GCF_000002035.6/53295/8090).

It should be noted that the latest versions of the nuclear genome assemblies of *Oryzias latipes* Hd-rR, *O. sakaizumii* HNI, *O. latipes* HSOK (as of April 2025) includes 896, 1453, and 1166 unanchored contigs that are not integrated into chromosomes, respectively. However, the genome assemblies that do not include these unanchored contigs remains as ‘references’ in NCBI. It is anticipated that genome sequences will be improved to achieve the ‘telomere-to-telomere (T2T)’ grade, as demonstrated recently in humans.^35^ In addition, numerous haploid genome assemblies were built for the Medaka Inbred Kiyosu-Karlsruhe (MIKK) panel, the first near-isogenic panel of 80 inbred lines in a vertebrate model derived from a wild founder population,^7^ with long reads with Oxford Nanopore Technologies but remain as non-chromosomal scaffolds.^36^

More recently, genome assemblies of wild relatives of Japanese medaka, such as the *Oryzias latipes* complex derived from East Asia^32^ and the Adrianichtydae species endemic to Sulawesi, Indonesia,^37^ has provided robust genomic resources for evolutionary and comparative biology^38^ (Figure 3). As of April 2025, NCBI Genome, the central archive for whole genome sequences covering the entire taxonomy, hosts genome assemblies for eight other species in the family Adrianichtyidae, namely *Adrianichthys oophorus*,^39^ *O. celebensis*,^37^ *O. dopingdopingensis, O. curvinotus*,^40^ *O*.*javanicus*,^41^ *O. woworae*,^37^ *O. sinensis*,^42^ and *O*.*melastigma*^43,44^ (recently recognized as *O. dancena* in Figure 1), as well as the two species (*O. luzonensis* and *O. mekongensis*) whose first genome assemblies are reported in this article (see below). The continuity and completeness of these genome resources largely vary depending mainly on the methodologies employed (Figure 3), which are expected to increase in upcoming years to cover a wider taxonomic range.

**Figure 3.**
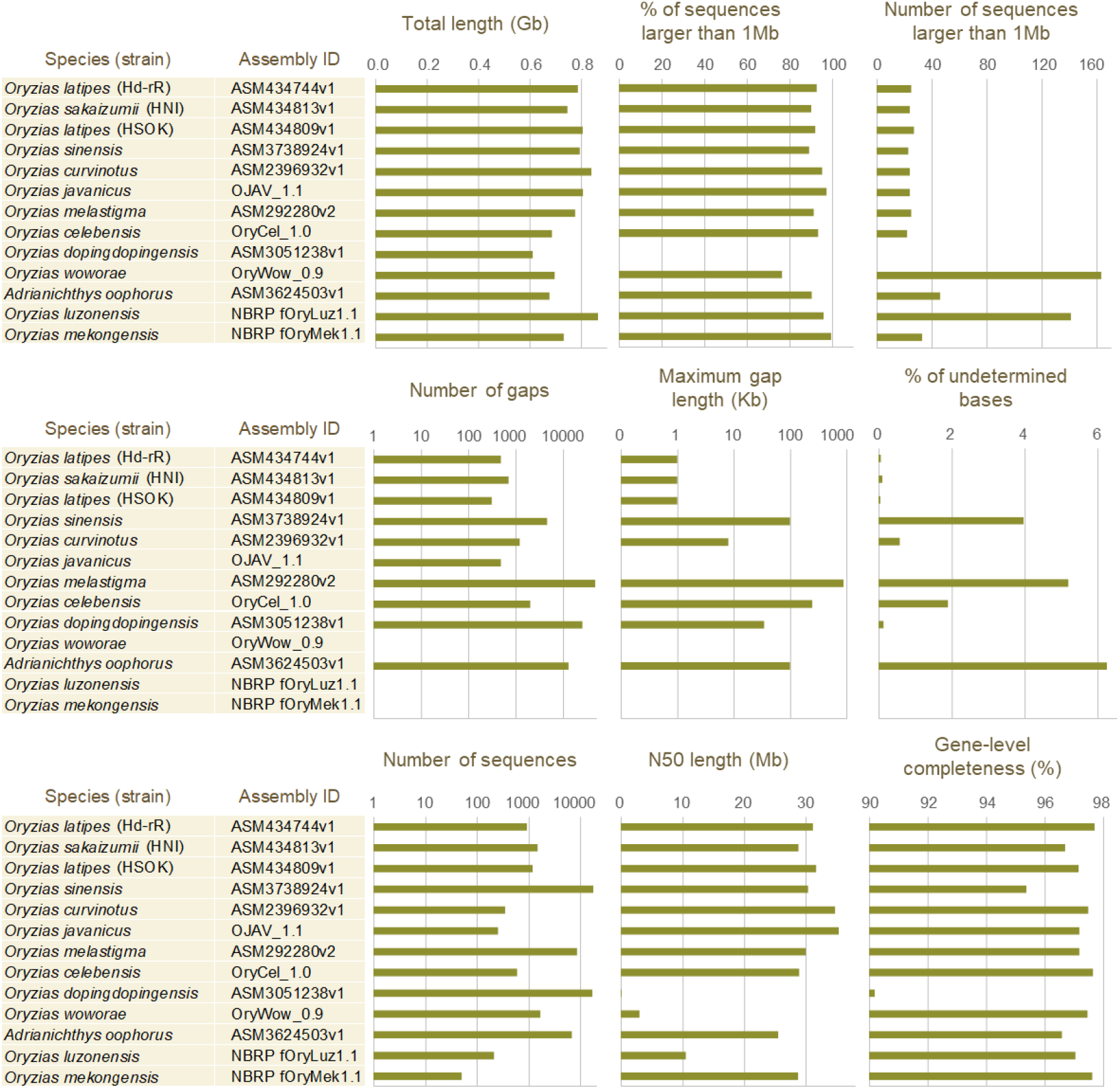
Statistics of the medaka genome assemblies. ‘Gaps’ denote those with no shorter than five undetermined bases. Gene-level completeness was evaluated by BUSCO v5.1.2^55^ on gVolante^54^ with the ortholog dataset Actinopterygii_odb10. See the ref. 56 for comparisons with more diverse teleost fish species. For Hd-rR, HNI, and HSOK, genome assemblies including unanchored contigs, whose Assembly IDs are different from those in the main text, are adopted for this statistics comparison.

### 3.2. Versatile web server: MedakaBase

The primary mission of NBRP Medaka’s activities is to facilitate the sharing of existing genomic resources in public databases and to promote their effective utilization. To this end, the MedakaBase service (Figure 4) was initiated at the end of 2022. The latest information about MedakaBase is announced in Medaka Omics News, and administrative information about server operation, such as scheduled downtime for maintenance, is communicated through MedakaBase X account (https://x.com/nbrpmedakaomix).

**Figure 4.**
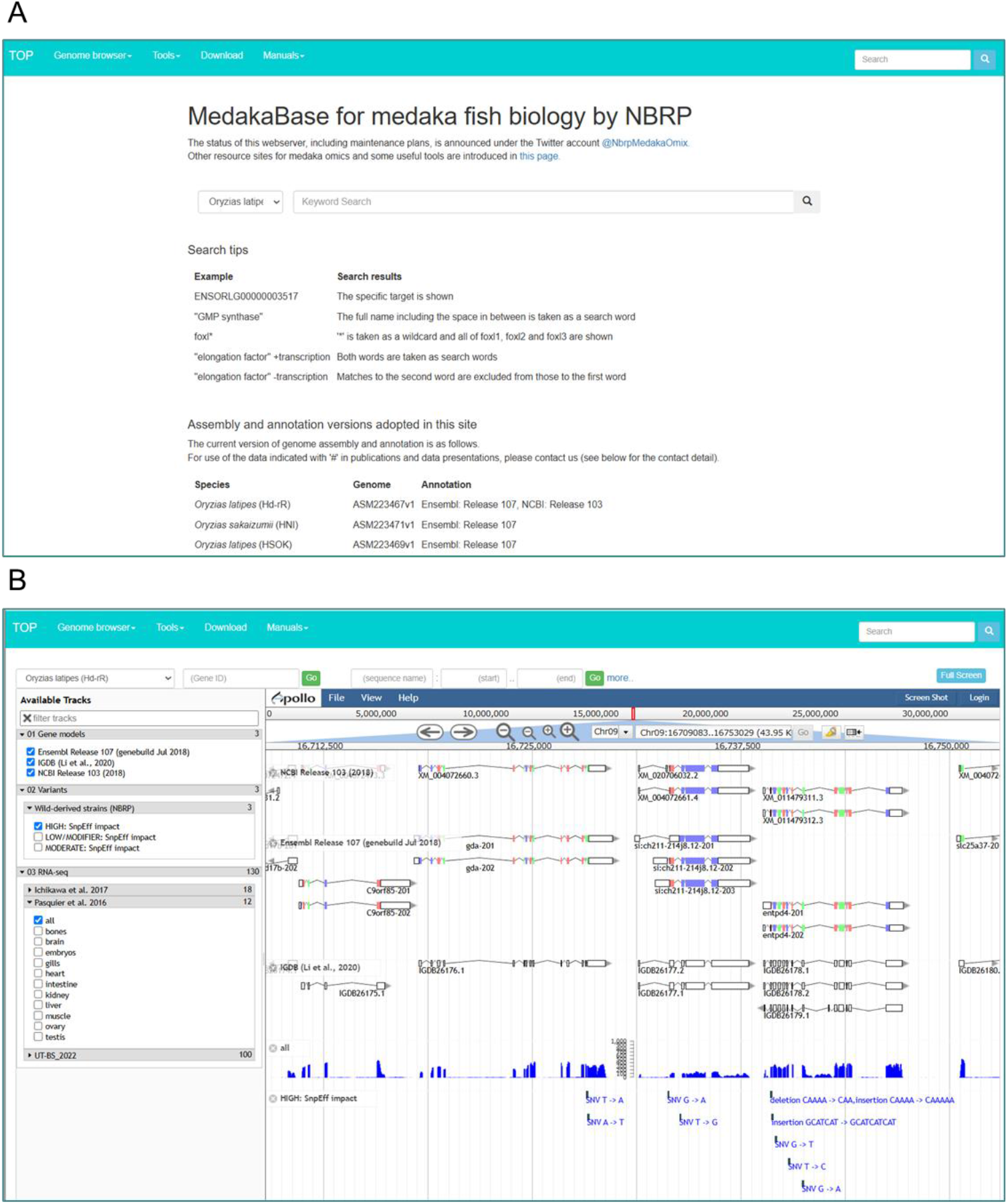
Web interface of MedakaBase. A, Top page. B, Genome browser that allows the choice of various genome-wide data (gene models, transcriptome, and SNPs) to visualize them in additional tracks in JBrowse^21^.

In the ‘JBrowse’ page (Figure 4B), users can display items of own interest from the checkboxes listed on the left. From the left lists, gene models can be selected from Ensembl, IGDB^45^ and/or NCBI, while RNA-seq data available in public^32,46,47^ can be displayed as bigWig files. Furthermore, modified GTF files and a modified GFF file by extending UTRs for advanced molecular analysis including single-cell analyses (see below), and the resultant files are now publicly accessible (https://figshare.com/projects/NBRP-Medaka/176391).

The ‘BLAST’ page of MedakaBase allows sequence similarity searches in publicly available and original genome-wide *Oryzias* sequence datasets. A search employs NCBI BLAST, using a single query input in a nucleotide or amino acid sequence. In MedakaBase, the abovementioned ‘unanchored contig’ sequences are included in the search targets of [Genome] in the Nucleotide databases. When hits are found in BLAST searches, they are explicitly labeled as “ unanchored contig”, distinct from chromosomal sequences.

In MedakaBase, the function ‘GMAP’ performs nucleotide sequence alignment of a given transcript onto the genome^48^, which allows the recognition of intronic sequences as unaligned regions between aligned regions. This facilitates the identification of exon ends, which is sometimes necessary for experimental design. Specifically, by selecting “ only show alignment” at the bottom of the [Format] popup menu, one can accurately determine, at a single basepair level, where introns are inserted into the cDNA sequence. Although the intron sequences are not displayed in this data, the insertion positions of introns are indicated by numbers. Using these numbers, the nucleotide sequence can be extracted with the Genome Slicer (see below). This can be applied to easily design gRNA on introns, 5′ UTRs, or 3′ UTRs. The ‘Genome Slicer’ function allows you to excise and display a specific region of a genome sequence based on specified coordinates for a given sequence ID (chromosome number). The ‘Gene Fetcher’ function enables the extraction of specific regions of a gene transcript by providing an Ensembl Transcript ID. From the top page of MedakaBase or by using the search window on the right side of the menu bar, users can perform a word search for genes. The search is conducted against the properties assigned to Ensembl gene entries. Information from Ensembl entries is based on Ensembl Release 107 (July 2022; gene build May 2018). Gene models starting with “ liftoff_” (available for HNI and HSOK) were generated using Liftoff ^49^ by the NBRP Medaka Team. It should be noted that they are based on cross-species inferences and are not supported by endogenous evidence for individual species.

These functions are available for five *Oryzias* genome assemblies (*O. latipes* Hd-rR and HNI [*O. sakaizumii*], HSOK, *O. cerebensis*, and *O. javanicus*) as of April 2025. The information for Ensembl entries available at MedakaBase is and will be based on Ensembl Release 107 (July 2022; gene build May 2018) unless stated otherwise.

### 3.3. UTR-aware gene models for single-cell RNA-seq

Additionally, we modified existing gene models to extend exons to cover 3′untranslated region (UTR) (Table 1). This process is demanded by possible data loss caused by the biased distribution of the reads obtained in some transcriptome sequencing methods including the Chromium Single Cell Gene Expression of 10X Genomics (Figure 5). This demand has also been evoked recently by studies focusing on mammals.^50^ For medaka Hd-rR, we modified existing GTF/GFF files by extending 3′-ends of transcripts using the program peaks2utr,^27^ and the technical note from our experience is publicly available at https://github.com/Squalomix/utr-modeling. Our UTR extension effort provided 3′ UTR annotation for 6,725 and 158 genes, for Ensembl (compared with v1) and NCBI (compared with v3), respectively, and achieved a higher mapping rate for the genomic regions recognized as exons (Table 1). The gene models of the Japanese medaka (*Oryzias latipes* Hd-rR) we modified are available at Figshare (https://figshare.com/projects/NBRP-Medaka/176391; DOI 10.6084/m9.figshare.24080463).

**Figure 5.**
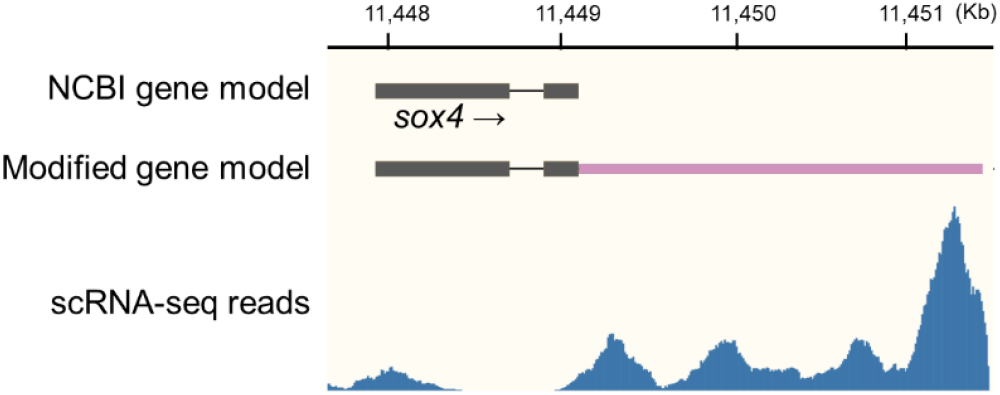
UTR extension for single-cell RNA-seq data analysis. Typical single-cell RNA-seq read mapping results (blue; see Methods) in comparison with the location of protein-coding exons (black) and the extended UTR (rose) for the *sox4* gene on chromosome 16 from the base position 11,447,921 to 11,449,123. The NCBI gene models and our modified gene models shown here were derived from NCBI GCF_002234675.1 and NBRP version 3 gene models, respectively.

### 3.4. Genome sequencing of other *Oryzias* species

To enrich genome resources of medaka fishes, we produced whole genome assemblies of *O. mekongensis* and *O. luzonensis* using animals supplied in the framework of NBRP. From a male of each species, we extracted high molecular weight DNA, processed it for single molecule, real-time (SMRT) sequencing and obtained circular consensus sequencing (CCS) (see Materials and Methods). The obtained high-fidelity long (HiFi) sequence data amounted to 38.2 and 25.6 Gb, respectively. Sequence assembly yielded 50 and 211 contigs for these species with N50 lengths of 28.7 and 10.5 Mb, respectively (Figure 3). These contigs exhibited percentages of detected one-to-one orthologs of over 97% suggesting a high completeness of protein-coding regions. Especially, the *O. mekongensis* genome contig sequences achieved a highest continuity, with more than 10 gapless chromosome-scale sequences flanked by conventional telomeric repeats (TTAGGG)n with no intervention by undetermined bases ‘N’.

For gene prediction on the obtained genome contigs, we employed a deep learning-based program Helixer^25^ that requires neither transcript evidence nor homolog hints. We obtained 28,683 and 29,544 protein-coding gene models for *O. mekongensis* and *O. luzonensis*, respectively. The obtained genome sequences and predicted gene sequences (both nucleotide and coding sequences) are available for BLAST searches at MedakaBase and for downloads at the dedicated FigShare archive linked from MedakaBase.

## 4. Discussion

MedakaBase provides a centralized platform for omics data of medaka fishes, integrating genome browsing, transcriptomic analyses, and comparative biology tools. Compared with previous databases, it enhances accessibility, data integration, and functionality for researchers. The platform enables BLAST-based searches, genome-wide exploration of variations, and user-friendly visualization. These features support evolutionary and developmental studies, particularly leveraging medaka as a model species. The curated datasets and streamlined interface improve the data retrieval and analysis. As different medaka fish species exhibit varying molecular mechanisms, e.g., in sex determination,^51^ functions enabling cross-species comparisons involving related species are expected to fuel medaka biology. Future updates will focus on expanding genomic data, such as T2T assembly, incorporating additional medaka strains, and integrating user-driven feedback. By continuously evolving under the framework of Life Science Infrastructure (LSI) renamed in 2025 from NBRP, MedakaBase aims to remain a critical resource for the research community, fostering discoveries in medaka genetics and beyond.

## Data availability

The genome assemblies reported in this article were registered under the BioProject PRJNA1240825 at NCBI.

## Acknowledgments

We acknowledge Mana Sato for cooperation in optimizing web server functions, Miwako Matsumoto and Hisayo Asao at NIBB for genomic HiFi read data production, animal caretakers for their daily effort for maintaining animals, Gaku Kimura for the maintenance of the web server MedakaBase, Masato Kinoshita, Yusuke Takehana, and Hideaki Takeuchi for discussion about web server usability and demands from biologists, Kazunori Yamahira, Soma Tomihara, Masaru Matsuda, Hiroyuki Takeda, and Shinichi Morishita for discussion about the utility of genome assembly and associated genome-wide resources. Computations were partially performed on the NIG supercomputer at ROIS National Institute of Genetics.

## Notes

https://figshare.com/projects/NBRP-Medaka/176391

### Competing Interest Statement

The authors have declared no competing interest.

## References

1. Parenti, L R., 2008, A phylogenetic analysis and taxonomic revision of ricefishes, Oryzias and relatives (Beloniformes, Adrianichthyidae), Zool. J. Linn. Soc., 154, 494–610.

2. Murata, K., Kinoshita, M., Naruse, K., Tanaka, M., and Kamei, Y. (eds.). 2019, Medaka: Biology, Management, and Experimental Protocols, John Wiley & Sons Ltd., Hoboken

3. Ishikawa, Y. 2000, Medakafish as a model system for vertebrate developmental genetics, Bioessays, 22, 487–495.

4. Wittbrodt, J., Shima, A., and Schartl, M. 2002, Medaka -a model organism from the far East, Nat. Rev. Genet., 3, 53–64.

5. Shima, A., and Mitani, H. 2004, Medaka as a research organism: past, present and future, Mech. Dev., 121, 599–604.

6. Takeda, H., and Shimada, A. 2010, The Art of Medaka Genetics and Genomics: What Makes Them So Unique?, Annu. Rev. Genet., 44, 217–241.

7. Fitzgerald, T., Brettell, I., Leger, A., et al. 2022, The Medaka Inbred Kiyosu-Karlsruhe (MIKK) panel, Genome Biol., 23, 59.

8. Aida, T. 1921, On the inheritance of color in a fresh-water fish, APLOCHEILUS LATIPES temmick and schlegel, with special reference to sex-linked inheritance. Genetics, 6, 554–573.

9. Yamamoto, T., 1958, Artificial induction of functional sex-reversal in genotypic females of the medaka (Oryzias latipes), J. Exp. Zool., 123, 571–594.

10. Matsuda, M., Nagahama, Y., Shinomiya, A., et al. 2002, DMY is a Y-specific DM-domain gene required for male development in the medaka fish, Nature, 417, 559–563.

11. Patyna, P. J., Davi, R. A., Parkerton, T. F., Brown, R. P., and Cooper, K. R. 1999, A proposed multigeneration protocol for Japanese medaka (Oryzias latipes) to evaluate effects of endocrine disruptors, Sci. Total Environ., 233, 211–220.

12. Ansai, S., and Kinoshita, M. 2014, Targeted mutagenesis using CRISPR/Cas system in medaka, Biol. Open., 3, 362–371.

13. Furutani-Seiki, M., Sasado, T., Morinaga, C.; et al. 2004, A systematic genome-wide screen for mutations affecting organogenesis in Medaka, Oryzias latipes. Mech. Dev., 121, 647–658.

14. Morinaga, C., Saito, D., Nakamura, S., et al. 2007, The hotei mutation of medaka in the anti-Mullerian hormone receptor causes the dysregulation of germ cell and sexual development, Proc. Natl Acad. Sci. USA, 104, 9691–9696.

15. Yokoi, H., Shimada, A., Carl, M., et al. 2007, Mutant analyses reveal different functions of fgfr1 in medaka and zebrafish despite conserved ligand-receptor relationships, Dev. Biol., 304, 326–337.

16. Montgomery, S. A., Tanizawa, Y., Galik, B. 2020, Chromatin Organization in Early Land Plants Reveals an Ancestral Association between H3K27me3, Transposons, and Constitutive Heterochromatin, Curr. Biol., 30, 573–588.

17. Tanizawa, Y., Mochizuki, T., Yagura, M., et al. 2025, MarpolBase: Genome database for Marchantia polymorpha featuring high quality reference genome sequences, bioRxiv, doi:10.1101/2025.03.30.646155

18. Matsumoto, Y., Chung, C. YL., Isobe, S. et al. 2024, Chromosome-scale assembly with improved annotation provides insights into breed-wide genomic structure and diversity in domestic cats, J. Adv. Res., in press.

19. Dunn, N. A., Unni, D. R., Diesh, C., et al. 2019, Apollo: Democratizing genome annotation, PLoS Comput. Biol., 15, e1006790

20. Priyam, A., Woodcroft, B. J., Rai, V., et al. 2019, Sequenceserver: A Modern Graphical User Interface for Custom BLAST Databases, Mol. Biol. Evol., 36, 2922–2924.

21. Diesh, C., Stevens, G. J., Xie, P., et al. 2023, JBrowse 2: a modular genome browser with views of synteny and structural variation, Genome Biol., 24, 74.

22. Altschul, S. F., Gish, W., Miller, W., Myers, E. W., and Lipman, D. J. 1990, Basic local alignment search tool, J. Mol. Biol., 215, 403–410.

23. Sim, S. B., Corpuz, R. L., Simmonds, T. J., and Geib, S. M. 2022, HiFiAdapterFilt, a memory efficient read processing pipeline, prevents occurrence of adapter sequence in PacBio HiFi reads and their negative impacts on genome assembly, BMC Genomics, 23, 157–163.

24. Cheng, H., Asri, M., Lucas, J., Koren, S., and Li, H. 2024, Scalable telomere-to-telomere assembly for diploid and polyploid genomes with double graph, Nat. Methods, 21, 967–970.

25. Holst, F., Bolger, A., Guenther, C., et al. 2023, Helixer–de novo Prediction of Primary Eukaryotic Gene Models Combining Deep Learning and a Hidden Markov Model, bioRxiv, doi: 10.1101/2023.02.06.527280

26. Kim, D., Paggi, J. M., Park, C., Bennett, C., and Salzberg, S. L. 2019, Graph-based genome alignment and genotyping with HISAT2 and HISAT-genotype, Nat. Biotechnol., 37, 907–915

27. Haese-Hill, W., Crouch, K. and Otto, T. D. 2023, peaks2utr: a robust Python tool for the annotation of 3’UTRs, Bioinformatics, 39, btad112.

28. Siddique, K., Ager-Wick, E., Fontaine, R., Weltzien, FA., and Henkel, C. V. 2021, Characterization of hormone-producing cell types in the teleost pituitary gland using single-cell RNA-seq, Sci. Data, 8, 279

29. Kasahara, M., Naruse, K., Sasaki, S., et al. 2007, The medaka draft genome and insights into vertebrate genome evolution, Nature, 447, 714–719.

30. Takeda, H., 2008, Draft genome of the medaka fish: A comprehensive resource for medaka developmental genetics and vertebrate evolutionary biology, Develop. Growth Differ., 50, S157–S166.

31. Kobayashi, D., and Takeda, H. 2008, Medaka genome project, Brief Funct. Genomic. Proteomic., 7, 415–426.

32. Ichikawa, K., Tomioka, S., Suzuki, Y., et al. 2017, Centromere evolution and CpG methylation during vertebrate speciation, Nat. Commun., 8, 1833.

33. Dyer, S. C., Austine-Orimoloye, O., Azov, A. G., et al. 2025, Ensembl 2025, Nucleic Acids Research, 53, D948–D957.

34. Bornstein, K., Gryan, G., Chang, E. S., Marchler-Bauer, A., and Schneider, V. A. 2023, The NIH Comparative Genomics Resource: addressing the promises and challenges of comparative genomics on human health, BMC Genomics, 24, 575.

35. Nurk, S., Kore, S., Rhie, A., et al. 2022, The complete sequence of a human genome, Science, 376, 44–53.

36. Leger, A., Brettell, I., Monahan, J., et al. 2022, Genomic variations and epigenomic landscape of the Medaka Inbred Kiyosu-Karlsruhe (MIKK) panel, Genome Biol., 23, 58

37. Ansai, S., Mochida, K., Fujimoto, S., et al. 2021, Genome editing reveals fitness effects of a gene for sexual dichromatism in Sulawesian fishes, Nat. Commun., 12, 1350.

38. Hilgers, L., and Schwarzer, J. 2019, The Natural History of Model Organisms: The untapped potential of medaka and its wild relatives, eLife, 8, e46994.

39. Yamahira, K., Kobayashi, H., Kakioka, R., et al. 2023, Ghost introgression in ricefishes of the genus Adrianichthys in an ancient Wallacean lake, J. Evol. Biol., 36, 1484–1493.

40. Li, M., Deng, A., He, C., et al. 2024, Genome sequencing, comparative analysis, and gene expression responses of cytochrome P450 genes in Oryzias curvinotus provide insights into environmental adaptation, Ecol. Evol., 14, e11565.

41. Takehana, Y., Zahm, M., Cabau, C. et al. 2020, Genome Sequence of the Euryhaline Javafish Medaka, Oryzias javanicus: A Small Aquarium Fish Model for Studies on Adaptation to Salinity, G3 (Bethesda), 10, 907–915.

42. Dong, Z., Wang, J., Chen, G., et al. 2024, A high-quality chromosome-level genome assembly of the Chinese medaka Oryzias sinensis, Sci. Data, 11, 322.

43. Lee, B-Y., Min-Sub Kim, M-S., Choi, B-S., et al. 2019, Construction of High-Resolution RAD-Seq Based Linkage Map, Anchoring Reference Genome, and QTL Mapping of the Sex Chromosome in the Marine Medaka Oryzias melastigma, G3 (Bethesda), 9, 3537–3545.

44. Liang, P., Saqib, H. S. A., Ni, X., and Shen, Y. 2020, Long-read sequencing and de novo genome assembly of marine medaka (Oryzias melastigma), BMC Genomics, 21, 640.

45. Li, Y., Liu, Y., Yang, H., Zhang, T., Naruse, K., and Tu, Q. 2020, Dynamic transcriptional and chromatin accessibility landscape of medaka embryogenesis, Genome Res., 30, 924–937

46. Pasquier, J., Cabau, C., Nguyen, T., et al. 2016, Gene evolution and gene expression after whole genome duplication in fish: the PhyloFish database, BMC Genomics, 17, 368.

47. Uchida, Y., Shigenobu, S., Takeda, H., Furusawa, C., and Irie, N. 2022, Potential contribution of intrinsic developmental stability toward body plan conservation, BMC Biol., 20, 8

48. Wu, T. D., and Watanabe, C. K. 2005, GMAP: a genomic mapping and alignment program for mRNA and EST sequences, Bioinformatics, 21, 1859–1875.

49. Shumate, A., and Salzberg, S. L. 2021, Liftoff: accurate mapping of gene annotations, Bioinformatics, 37, 1639–1643.

50. Pool, AH., Poldsam, H., Chen, S., Thomson, M., and Oka, Y. 2023, Recovery of missing single-cell RNA-sequencing data with optimized transcriptomic references, Nature Methods, 20, 1506–1515.

51. Takehana, Y., Matsuda, M., Taijun Myosho., et al. 2014, Co-option of Sox3 as the male-determining factor on the Y chromosome in the fish Oryzias dancena, Nat. Commun., 5, 4157

52. Yamahira, K., Ansai, S., Kakioka, R., et.al. 2021, Mesozoic origin and ‘out-of-India’ radiation of ricefishes (Adrianichthyidae), Biol. Lett., 17, 20210212.

53. Sayers, E.W., Cavanaugh, M., Frisse, L., et al. 2025, GenBank 2025 update, Nucleic acids research, 53, D56–D61.

54. Nishimura, O., Hara, Y. and Kuraku, S. 2017, gVolante for standardizing completeness assessment of genome and transcriptome assemblies, Bioinformatics, 33, 3635–3637.

55. Seppey, M., Manni, M., and Zdobnov, E. M. 2019, BUSCO: Assessing Genome Assembly and Annotation Completeness, Methods in Molecular Biology, 1962, 227–245

56. Kuraku, S., Sato, M., Yoshida, K., and Uno, Y., 2024, Genomic reconsideration of fish non-monophyly: why cannot we simply call them all ‘fish’?, Ichthyol. Res., 71, 1–12

